# A prediction-interval framework for identifying outlier control cohorts in lifespan studies

**DOI:** 10.64898/2026.10.21.760720

**Authors:** Laurence T. Maeyens, Alison Luciano, James F. Nelson, Gary A. Churchill

## Abstract

Control cohorts anchor the interpretation of lifespan-extending interventions, yet their lifespans vary even under identical experimental conditions. Without quantifying this expected variation, it is difficult to distinguish unusually short- or long-lived cohorts arising by chance from those reflecting potential experimental anomalies, with important consequences for interpreting treatment effects. The National Institute on Aging Interventions Testing Program (ITP), which tests interventions in mice across three independent sites using standardized protocols, provides a unique opportunity to address this problem. Previously, unusually short-lived males at the University of Texas in 2017 and unusually long-lived females at The Jackson Laboratory in 2022 were identified as potentially anomalous, but no standardized framework existed to determine whether such deviations exceeded expected variation. Here, we analyzed 10,757 control mice spanning 18 years of the ITP to quantify historical variation within and between sites and develop a prediction-interval framework for prospectively identifying unusual cohorts. We found that year-to-year variation, particularly in males, contributed substantially more to variation among cohort means than expected from random sampling alone. Accounting for this variation, our predictive framework flagged the 2022 Jackson Laboratory females but not 2017 University of Texas males, while flagging a previously unrecognized male cohort. Excluding this newly flagged cohort from the original lifespan analysis changed the statistical significance of two rapamycin regimens. These findings highlight the importance of a standardized framework for outlier detection and demonstrate the importance of accounting for natural variability in aging studies.

## Introduction

Reproducibility is a foundational concern in aging research, where subtle intervention effects and long experimental timelines make it difficult to distinguish a genuine treatment effect from noise. Interlaboratory testing, in which the same protocol is run in parallel at independent sites, is one of the most direct ways to guard against site-specific artifacts, but it depends on researchers being able to distinguish genuinely aberrant variation from ordinary variability^1;2^.

Lifespan studies remain the most direct way to test whether an intervention alters the trajectory of aging and survival. The National Institute on Aging Interventions Testing Program (ITP) is one of the most rigorous platforms for testing candidate lifespan-extending interventions in mice. The program was designed to improve reproducibility in aging intervention studies by testing compounds in parallel across three independent sites using genetically heterogeneous HET3 mice of both sexes, standardized protocols, adequate sample sizes, and parallel control cohorts^3–7^. Because interventions have been evaluated against parallel control cohorts for 18 completed years, spanning the 2004 through 2022 cohorts, the program provides an unusually rich historical dataset for studying lifespan variability, comprising lifespan data from over 10,000 control animals.

However, lifespan is also an inherently variable outcome. Even among genetically identical animals kept under the same controlled conditions, lifespans can differ substantially, reflecting the inherently stochastic nature of aging, disease development, and mortality. Despite the rigor of the ITP, control cohorts are also not identical from year to year. This is not unexpected: in addition to intrinsic individual-level variation in lifespan, subtle differences in husbandry, environmental conditions, microbiological status, or experimental implementation across sites and over time may influence lifespan ^8^. This creates a practical problem for interpreting intervention studies: if a control cohort has an unusually short or unusually long lifespan, every treatment tested against that control cohort may appear weaker or stronger than they are on average.

Some cohorts may appear visually unusual, raising the question of whether they reflect ordinary stochastic variation or a true procedural, environmental, or biological anomaly. This distinction matters because an unusually short-lived control group could exaggerate apparent treatment benefits, whereas an unusually long-lived control group could obscure a real intervention effect. The cost runs in both directions. Discarding a control cohort that was in fact ordinary biases control estimates, which in turn biases intervention efficacy estimates for every intervention tested at that site in that year. If different rules are used to discard purported controls in different years, the extent of bias is variable, breaking year-to-year comparability. Hierarchical models that explicitly partition site- and year-level variability from treatment effects offer one route to making this distinction rigorous ^9;10^. Yet there is currently no widely adopted statistical framework for deciding when a control cohort is sufficiently unusual to warrant further investigation^11;12^.

We present two examples from the National Institute on Aging Interventions Testing Program to highlight the need for such a framework. In the 2017 ITP cohort, control mice at a single site appeared unusually short-lived by median lifespan, particularly in males^13^. Conversely, in the 2022 ITP cohort, female control mice at a single site appeared unusually long-lived, driven in part by several animals surviving to very old ages^14^. A prespecified predictive framework is needed to determine when a control cohort should be flagged for further investigation.

We approach this in three steps. Because the program has run the same protocol in parallel at three sites for eighteen years, the control record can be used to measure variability between laboratories and between years, rather than assuming it. We first characterize and partition historical variation into within-cohort, between-year, and between-site components; second, we construct historical reference ranges from the completed control record; and third, we use rolling cumulative data windows to ask prospectively whether a new cohort would have been flagged using only preceding controls.

## Results

The ITP control record is an opportunity to study interlaboratory reproducibility: the same protocol, run in parallel at three sites on the same genetic background, for eighteen years. We use it first to measure by how much control outcomes vary between laboratories and between years, and then to ask when a cohort has varied further than that historical record can explain.

### Interlaboratory variation in control lifespan across three sites and eighteen years

The National Institute on Aging Interventions Testing Program includes three testing sites: The Jackson Laboratory (TJL), the University of Michigan (UM), and the University of Texas Health Science Center at San Antonio (UT). Across these sites, the historical control data show that lifespan variability is influenced by both sex and site (Figure 1). Females lived longer than males on average at all three sites (Figure 1A and 1B), as previously reported ^15^. Male survival curves were broader and more gradual, with more pronounced early mortality and greater site-to-site separation (Figure 1B). These patterns support the overall conclusion that male control lifespan is less stable across cohorts and sites^5^. However, this sex difference was least pronounced at UM, where male controls had a substantially longer median lifespan (~10%) than males at the other two sites (Figure 1B). UM showed the greatest between-cohort variation in male lifespan, which reflects the fact that male controls at this site have often been nearly as long-lived as females, whereas from approximately 2015 to 2018 male lifespan was markedly shorter and more closely resembled the male–female separation observed at the other two sites. The lowest annual female Kaplan–Meier (KM) median lifespan was observed at UT in C2017 (827 days), whereas the highest was observed at TJL in C2022 (948 days). Among males, the extremes were 667 days at TJL in C2018 and 927 days at UM in C2010. A site-by-sex temporal analysis suggested an increase in annual Kaplan–Meier median lifespan among TJL females, although the association did not remain significant after correction for multiple testing (r = 0.55, P = 0.018, q = 0.110). The positive trend persisted but weakened after excluding C2022 (r = 0.46, P = 0.066), suggesting that the unusually high C2022 median may partly reflect a longer-term temporal increase rather than an isolated departure from historical patterns (Supplementary Figure 1). Two cohorts previously highlighted in ITP publications, C2017 UT males and C2022 TJL females ^13;14^, warranted formal analysis to determine whether their classification as outliers was statistically justified. Importantly, in any historical dataset there will always be a highest and lowest cohort value; the key question is whether these extremes are larger than expected under normal year-to-year variation. The 2022 TJL female cohort and the 2017 UT male cohort therefore provide useful case studies for formally testing whether visually extreme control cohorts should be considered statistical outliers or interpreted as part of expected historical variability.

**Figure 1.**
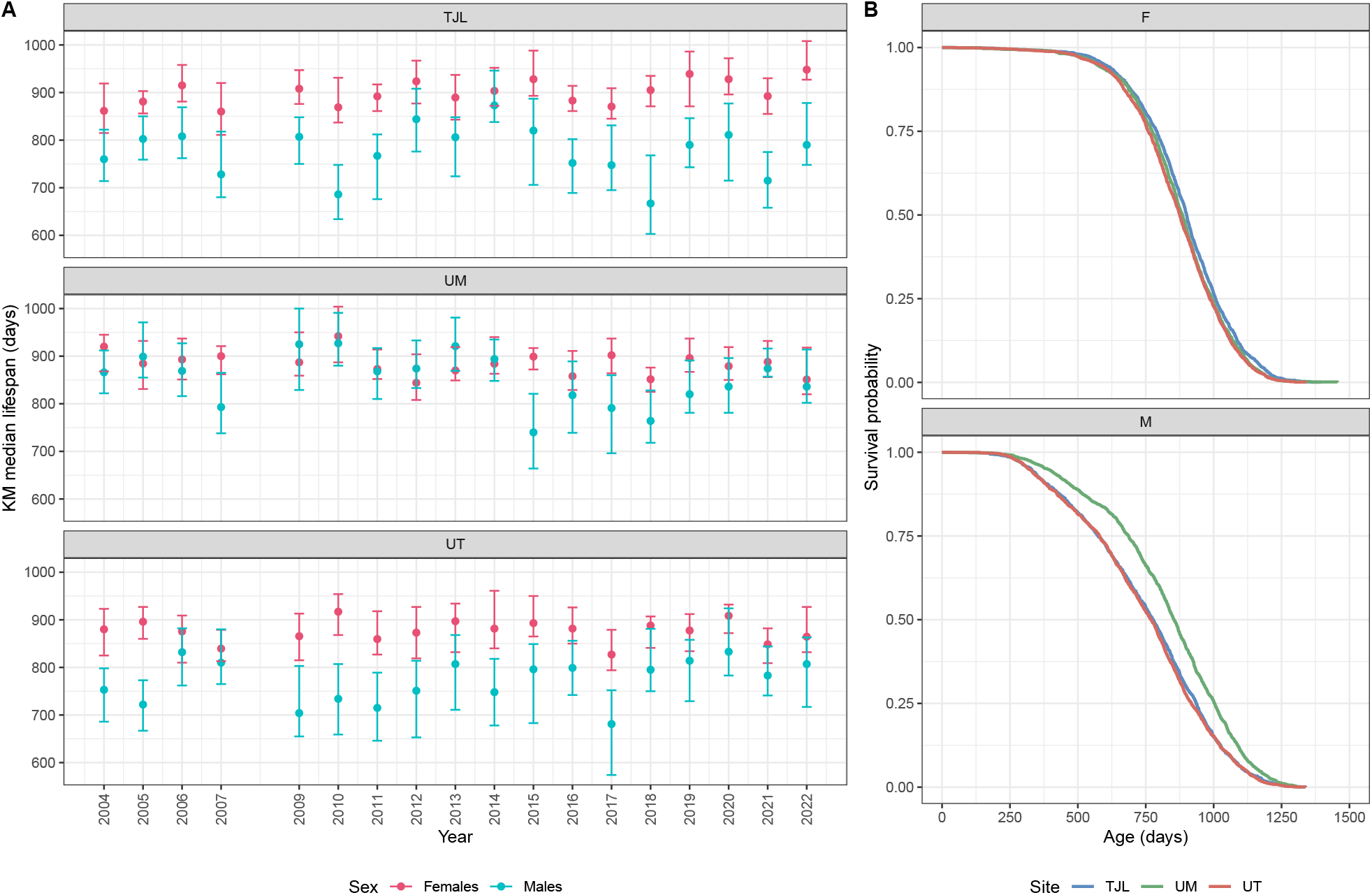
Historical variation in control lifespan across site, sex, and year in the National Institute on Aging Interventions Testing Program. A) Annual Kaplan–Meier median lifespan of control mice from 2004 to 2022, stratified by testing site and colored by sex. Points show site-by-sex cohort Kaplan–Meier medians, and error bars represent 95% confidence intervals. B) Kaplan–Meier survival curves pooling all years, shown separately for males and females and colored by site. Survival differed more across sites in males than in females, consistent with greater male lifespan variability.

**Table 1.** Summary of between-cohort variation across annual site-by-sex control cohorts. Mean observed age, between-cohort SD, range, and CV summarize annual cohort means; pooled Kaplan–Meier medians correspond to Figure 1B, while the accompanying SD and range summarize variation among annual cohort-specific Kaplan–Meier medians. Mean within-cohort SD reflects mouse-to-mouse variation within annual cohorts.

| Between-cohort variation in control lifespan |  |  |  |  |  |  |  |  |  |  |
| --- | --- | --- | --- | --- | --- | --- | --- | --- | --- | --- |
| Variation across annual site-by-sex control cohorts |  |  |  |  |  |  |  |  |  |  |
| Sex | Cohorts | Total N | Annual cohort means |  |  |  | Kaplan-Meier medians |  |  |  |
|  |  |  | Mean observed age | Between-cohort SD | Range | CV (%) | Pooled KM median | Between-cohort SD | Range | Mean within-cohort SD |
| TJL |  |  |  |  |  |  |  |  |  |  |
| F | 18 | 1,726 | 880.3 | 37.3 | 143.2 | 4.2 | 900.0 | 26.6 | 88.0 | 177.3 |
| M | 18 | 1,908 | 729.4 | 35.3 | 117.8 | 4.8 | 776.0 | 53.6 | 206.0 | 249.9 |
| UM |  |  |  |  |  |  |  |  |  |  |
| F | 18 | 1,671 | 865.9 | 22.6 | 91.7 | 2.6 | 885.0 | 24.8 | 98.0 | 185.0 |
| M | 18 | 1,880 | 766.5 | 70.1 | 240.8 | 9.1 | 854.0 | 55.1 | 187.0 | 267.5 |
| UT |  |  |  |  |  |  |  |  |  |  |
| F | 18 | 1,708 | 859.8 | 22.3 | 100.1 | 2.6 | 875.0 | 23.1 | 90.0 | 180.9 |
| M | 18 | 1,864 | 734.6 | 47.7 | 181.8 | 6.5 | 771.0 | 46.3 | 152.0 | 240.4 |
| Annual means and within-cohort SDs use observed ages; pooled and annual medians are Kaplan-Meier estimates. |  |  |  |  |  |  |  |  |  |  |

Annual means and within-cohort SDs use observed ages; pooled and annual medians are Kaplan–Meier estimates.

### Partitioning control-cohort variation into within-cohort, between-year and between-site components

Cohort-level summaries describe by how much control outcomes vary but not where that variation comes from. To separate sources of variance, we modeled observed age at the level of the individual mouse, with sex *×* site fixed effects and Site *×* Year random effects. Because female and male cohorts showed different degrees of annual variability, the random-effect structure allowed separate Site *×* Year variances by sex. A model allowing separate female and male Site *×* Year variances was strongly favored over a model imposing a single shared Site *×* Year variance across sexes (ΔAIC = 50.9; ΔBIC = 43.6). Residual variance was also allowed to differ by sex, which was strongly supported compared with a homogeneous-variance model (likelihood-ratio test, *χ*^2^(1) = 543.5, P = 3.29 *×* 10^*−*120^). Interval estimates were obtained by wild bootstrap (Methods).

At the individual-mouse level, residual variation remained substantial, with residual standard deviations of 183.2 days in females and 253.4 days in males. However, this variation is averaged across animals when considering a cohort mean, corresponding to sampling standard errors of 18.8 days for females and 24.8 days for males at mean cohort sizes of 94.5 and 104.7 animals, respectively. Allowing Site*×* Year variance to differ by sex revealed appreciable year-to-year variation, with Site *×* Year standard deviations of 21.1 days in females and 46.7 days in males. Systematic differences among the three fitted site means were comparatively modest, with 20.4 days separating the highest and lowest female estimates and 37.2 days separating the corresponding male estimates.

Expressed as proportions of the estimated variance in an observed cohort mean, between-year variation was the largest component, accounting for 50.9% of the variance in females and 71.1% in males. Individual-level sampling variation accounted for 40.7% and 20.0%, respectively, whereas systematic differences among the three ITP sites accounted for 8.4% and 8.9%. The resulting estimated standard deviation of an observed cohort mean was 29.5 days in females and 55.4 days in males. Thus, male control cohorts were not only more variable at the individual level but also exhibited substantially greater year-to-year variability, which became the dominant source of variation in male cohort means. Male cohort means are less stable because most of their variation comes from site-by-year shifts, not sampling noise. Increasing cohort size reduces the sampling component, but it cannot remove genuine year-to-year changes in the control population.

**Table 2.** Sex *×* site lifespan estimates from the individual-level heterogeneous-variance model. Fixed effects included sex, site, and their interaction; Site *×* Year was modeled as a random effect with separate variance components for females and males, and residual variance was allowed to differ by sex. Lifespan (d) is estimated mean observed age in days; intervals are percentile intervals from 5,000 wild-bootstrap resamples (Mammen auxiliary distribution, HC2 weighting) of the fit to the complete record.

| Site | Sex | Lifespan (d) | 99% CI, wild bootstrap (d) |
| --- | --- | --- | --- |
| TJL | F | 880.2 | 859.2 – 901.2 |
| TJL | M | 729.5 | 709.0 – 748.6 |
| UM | F | 865.8 | 852.3 – 877.1 |
| UM | M | 766.7 | 724.7 – 802.8 |
| UT | F | 859.8 | 847.0 – 871.8 |
| UT | M | 734.4 | 706.1 – 761.7 |

We also examined variation in censoring frequency across sites, sexes, and cohort years (Supplementary Figure 8). Censoring was particularly frequent among UM males (12.2%), compared with TJL males (5.3%) and UT males (2.9%). Censoring frequency also varied considerably between cohort years, and censored animals generally had younger recorded ages than animals recorded as natural deaths. The components above describe variation among control cohorts. Turning that description into a decision about a particular cohort requires a reference range, and a statement of what falling outside it does and does not establish.

### A reference range built from the complete control cohort record

A natural first approach is to build a reference range directly from the completed record, maximizing information use. We modeled the Kaplan–Meier median lifespan estimated for each site *×* sex *×* year cohort as a function of sex, with testing site as a random effect and separate residual variability for females and males, treating sites as exchangeable rather than deriving site-specific thresholds. Fitting to all completed cohorts from C2004–C2022 and simulating sex-specific predictive distributions gave a model-predicted center of 886.9 days for female cohorts (99% interval 816.3–957.0) and 799.5 days for male cohorts (99% interval 640.0–959.2; Figure 2). The 99% intervals contained all 54 female and all 54 male cohorts (Supplementary Figure 2). The two cohorts of interest, C2022 TJL females and C2017 UT males, fell within their respective sex-specific intervals. Excluding these target years from model training did not alter these median-based classifications. Adding a year trend, site × sex interaction, site- and sex-specific temporal drift, or cohort size likewise did not flag any historical cohort. Model fit was improved by the site *×* sex and, most strongly, the site- and sex-specific temporal-drift extensions, with the latter also improving fit relative to the site × sex model (likelihood-ratio test, P = 0.0192). Using cohort mean observed age rather than Kaplan–Meier median lifespan identified C2022 TJL females as exceeding the 99% prediction interval, whereas C2009 TJL females and C2017 UT males remained within their respective intervals. In contrast, using raw medians of observed age identified C2009 TJL females as the only cohort outside the 99% prediction interval. However, this cohort remained within the interval when Kaplan–Meier medians were used, even after excluding all C2009 cohorts from model fitting. These findings illustrate how outlier classifications can depend on the choice of lifespan summary.

**Figure 2.**
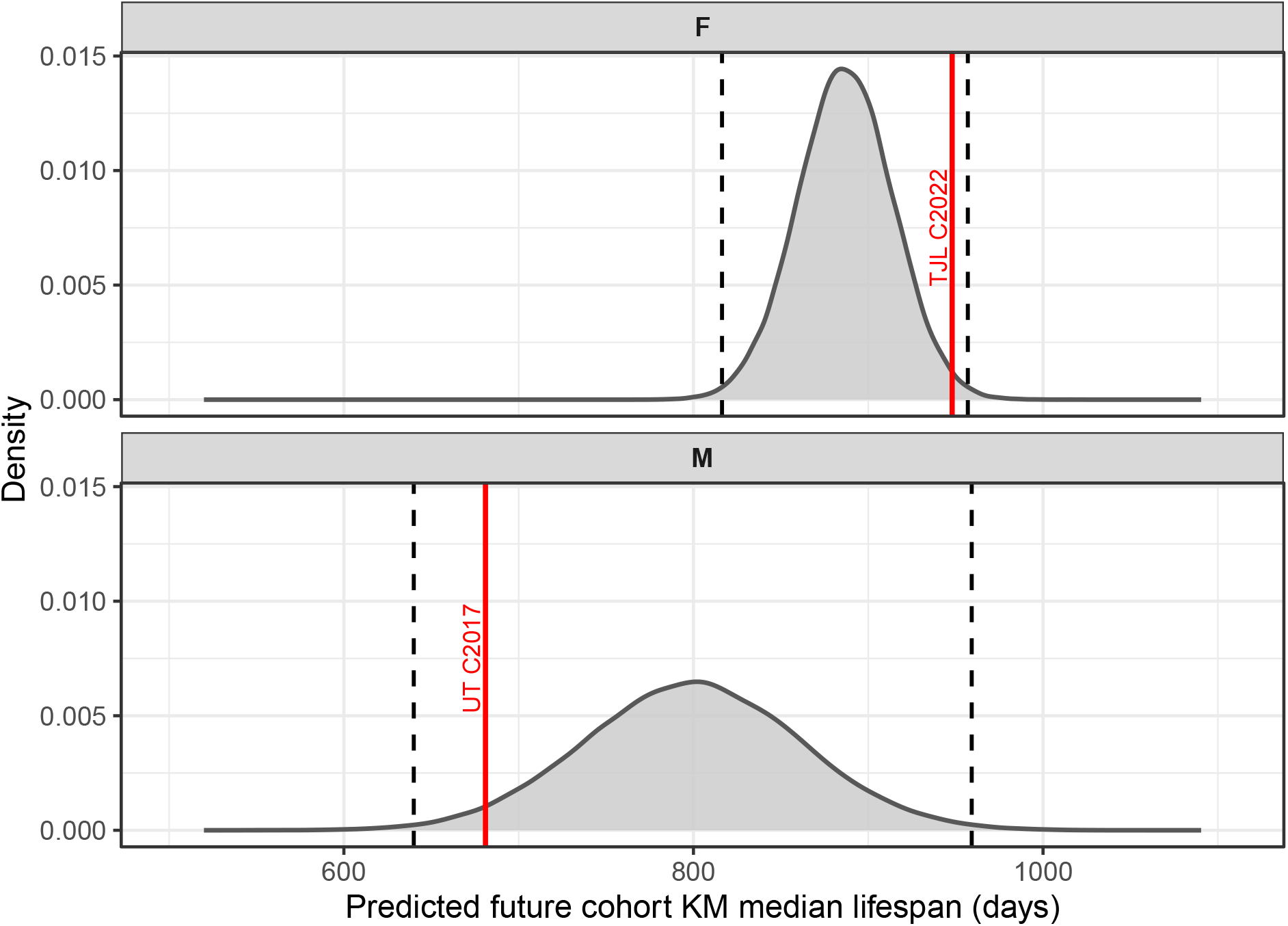
Site-exchangeable prediction intervals for historical control-cohort Kaplan–Meier median lifespan. Predictive distributions for the estimated Kaplan–Meier medians of female and male control cohorts were generated from the cohort-level mixed-effects model fitted to the complete record, treating testing sites as exchangeable. Grey density curves show simulated cohort Kaplan–Meier median estimates, and dashed vertical lines indicate the 99% prediction interval. Red lines indicate the observed Kaplan–Meier medians of C2022 TJL females and C2017 UT males.

The limitation is that this is an in-sample assessment: the same 108 cohorts were used to fit and evaluate the reference model. The observed historical coverage should therefore be viewed as descriptive rather than as independent validation. Under a correctly calibrated 99% predictive distribution, approximately 1% of future cohorts would be expected to fall outside the interval by chance. Therefore, roughly one of 108 cohorts would be expected to fall outside the interval by chance alone; even if a single in-sample exceedance had been observed, this would not, by itself, establish that the cohort was truly anomalous. Evaluating whether a cohort is statistically unusual requires applying a prespecified rule using only preceding data.

### Prospective detection of outlier control cohorts

A prospective question instead concerns a single cohort nominated in advance: e.g., this year’s UT males have finished, and we ask whether they fall within an expected range. For the prospective question, six control cohorts complete each year (3 sites *×* 2 sexes), each carrying a 1-in-100 chance of a false alarm (99% interval) under perfect calibration, so the rule raises about six false alarms per century, or one every seventeen years, and approximately one across the eighteen completed years of the program.

To assess how the framework would behave if applied in real time, we carried out an internal temporal validation using future years drawn from the same historical record. We refit the model and repeated the bootstrap on successive cumulative windows, beginning with the earliest cohort years and adding one year at a time. We then asked whether each window’s prediction interval would have contained the following year’s observed cohort mean. In other words, whether the current year’s observations would be considered outliers given the historical record available up to that point in time. For each window, prediction intervals incorporated uncertainty in the fitted sex *×* site mean together with the additional variability expected in a future cohort. Specifically, for each wild-bootstrap estimate of the site *×*sex mean, we simulated a future sex-specific site *×* year effect and cohort-mean residual variation based on the observed size of the subsequent cohort. The 0.5th and 99.5th percentiles of these simulated future cohort means defined the 99% prediction interval. Between-year variance therefore enters the prediction interval, whereas site variance does not because prediction is conditional on the known testing site but marginal over the future year. Estimates and their uncertainty were monitored across cumulative historical windows as additional cohort years became available (Supplementary Figure 5).

Because uncertainty estimates were unstable in short historical windows, we retained eight completed cohort years as a pragmatic minimum for the primary prospective analysis. The average ratio of bootstrap-based to model-based standard errors had largely stabilized by approximately seven to eight historical cohort years and showed little systematic improvement thereafter, although greater variability remained among individual site *×* sex estimates (Supplementary Figure 6). Among windows meeting the eight-year minimum, the nominal 99% prediction interval contained the subsequent cohort mean in 57 of 60 comparisons (95.0%; Figure 3). The three cohorts falling outside their prospectively generated intervals were UM males in C2015 and C2018 and TJL females in C2022. Thus, the revised model continued to flag the unusually long-lived C2022 TJL female cohort, whereas the C2017 UT male cohort fell within its prospectively generated interval once sex-specific year-to-year variability was incorporated. The latter result indicates that its unusually short observed lifespan was nevertheless compatible with the degree of male annual variation estimated from preceding control cohorts.

**Figure 3.**
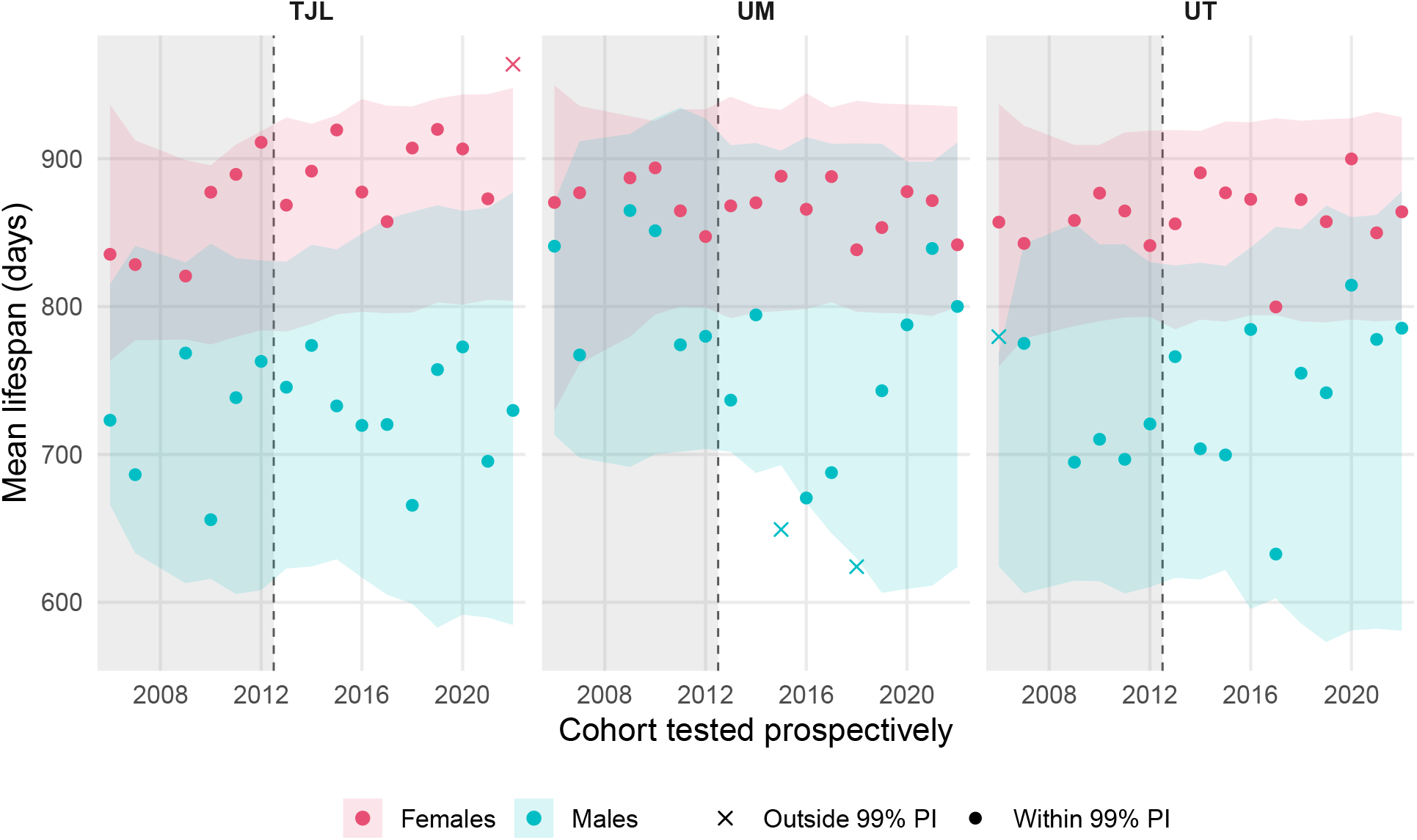
Prospective validation of the prediction framework. For each cohort shown on the x-axis, the model was fit using only control cohorts available before that cohort and used to generate a 99% prediction interval (shaded band) for its cohort’s mean observed age. Points indicate the observed mean lifespan of the cohort being tested. Panels indicate site, colors indicate sex, and point shape denotes whether the observed cohort mean fell within or outside its prediction interval. The gray region marks predictions based on fewer than the eight historical cohort years required for the primary analysis. Among the 60 comparisons meeting this threshold, 57 (95.0%) were contained within the nominal 99% prediction interval.

We specified a priori that temporal drift would be excluded from the primary prediction framework and evaluated only in a sensitivity analysis. Because the descriptive temporal analysis suggested potential site- and sex-specific longitudinal structure (Supplementary Figure 1), we additionally tested whether allowing a separate linear temporal slope for each site *×* sex combination improved prospective prediction. The sensitivity model included sex, site, centered cohort year, and all two- and three-way interactions, while retaining the primary model’s sex-specific Site *×* Year random effects and residual variances. Adding site- and sex-specific linear temporal drift did not improve prospective coverage, which remained 57 of 60 comparisons (95.0%) under both models, while the mean 99% prediction-interval width increased slightly from 192.5 to 195.1 days. The drift model contained the C2018 UM male and C2022 TJL female observations that were flagged by the base model but instead flagged C2017 UT females and barely C2021 UM males, the latter exceeding the upper 99% prediction limit by only 0.3 days. C2015 UM males remained the only cohort flagged under both specifications. Consistent with the a priori analysis plan, the model without temporal drift remained the primary framework; the sensitivity analysis showed that allowing drift changed cohort classifications without improving coverage or narrowing the prediction intervals.

Across the complete set of rolling windows, including those below the eight-year threshold, 92 of 96 subsequent cohorts were contained (95.8%), while 35 of 36 (97.2%) were contained among the shorter windows alone. With 60 primary comparisons, a nominal 99% interval would be expected to miss fewer than one cohort on average; the three observed misses may therefore reflect genuinely unusual cohorts, remaining unmodeled temporal structure, or some combination of the two. Cohorts falling outside the prediction interval should consequently be interpreted as statistical flags warranting further investigation rather than definitive outliers.

Because C2015 UM males were the only cohort flagged under both the primary and temporal-drift prospective models, and because the consequences of excluding this cohort had not previously been evaluated, we next asked whether it materially affected interpretation of the interventions tested in C2015. We repeated the male treatment analyses after excluding all UM males, including both control and treatment-assigned animals, while retaining the original site-stratified log-rank framework ^16^. Exclusion of UM increased the control median lifespan from 775 to 806 days. The survival benefit of Rapa 20 remained significant, although its relative median-lifespan effect decreased from 10.7% to 6.9% (P = 6.57 *×* 10^*−*4^ versus P = 0.0022). In contrast, Rapa cycles changed from a 9.3% increase in median lifespan (P = 0.0019) to a 3.6% decrease (P = 0.217), and Rapa 20–23 changed from a 10.7% increase (P = 0.024) to a 6.0% increase that was no longer significant (P = 0.085). These changes should be interpreted cautiously because excluding an entire testing site also reduces sample size and statistical power. Consequently, changes in statistical significance cannot be attributed solely to the removal of an unusual control cohort. The remaining interventions remained nonsignificant in both analyses (Supplementary Figure 7).

## Discussion

A single striking result, however unusual it looks, cannot on its own establish whether a control cohort is a true anomaly: rare events are expected to occur at their expected rate, however surprising any one occurrence feels, and no isolated observation can substitute for a reliable body of comparable evidence^17^. The same logic applies here. Our analysis provides a quantitative basis for flagging control cohorts that fall outside the range expected under the historical record for further investigation.

Framed more generally, this analysis treats the ITP control record as a long-running study in interlaboratory reproducibility. Three sites have run the same protocol in parallel on the same genetic background for eighteen years, which is an unusual opportunity in aging biology to measure, rather than assume, if and to what extent lifespan is variable in controls between laboratories and over time. We use this record in three complementary ways: first to characterize and partition variation among control cohorts, second to construct historical reference ranges, and third to ask prospectively whether a new cohort would have been flagged using only information available before that cohort occurred.

The distinction between retrospective and prospective assessment is important. A cohort identified by searching a completed historical record carries less evidential weight than one flagged by a rule specified using only preceding data, because the most extreme observation in a sufficiently large dataset will necessarily appear unusual. This distinction was apparent here. In the complete-record Kaplan–Meier median-lifespan analysis, none of the 108 historical cohorts fell outside the sex-specific 99% prediction intervals, including the two cohorts that motivated this analysis, C2017 UT males and C2022 TJL females. Under prospective evaluation using the individual-level model, C2022 TJL females were flagged by the primary model but not when site- and sex-specific temporal drift was included, indicating that their classification was sensitive to model specification. In contrast, C2017 UT males were not flagged under either specification.

Thus, our analysis weakens the statistical justification for treating C2017 UT males as an outlier^13^, whereas the evidence for C2022 TJL females receives partial support^14^. This distinction is consequential, as exclusion of UT in the C2017 study eliminated the significant pooled male captopril effect and the small female butanediol effect. In contrast, C2015 UM males were flagged under both the primary and temporal-drift models, making this the most consistently flagged cohort across the two prospective specifications. Although unusual site-specific survival patterns were noted in the original C2015 report^16^, the UM male control cohort was not formally evaluated against the historical control record or excluded in a secondary sensitivity analysis. Here, excluding UM males eliminated the significant survival effects of the cycling and 20–23-month rapamycin regimens, whereas the 20-month rapamycin regimen remained significant. This illustrates the value of applying a formal, prespecified assessment to each newly completed control cohort and using statistical flags to trigger sensitivity analyses rather than relying on visual inspection or ad hoc judgment. The prospective framework did not simply confirm visually extreme cohorts: it distinguished observations that remained unusual after accounting for historically estimated sex-specific year-to-year variability from those that could plausibly arise within that variation.

The variance partitioning also has design implications. After individual-level variation was averaged over cohort size, site *×* year variation was the largest contributor to control-cohort mean variability, accounting for 50.9% in females and 71.1% in males, whereas sampling variation accounted for 40.7% and 20.0%, respectively; differences among fitted site means contributed relatively little. Increasing the number of parallel controls can reduce sampling variation but cannot eliminate genuine year-to-year shifts in the control population. Importantly, however, site *×* year variation does not necessarily translate directly into poorer treatment-effect precision, because treatment and parallel control groups are tested under the same site-year conditions and shared shifts may cancel in the treatment–control contrast. It becomes more consequential when those shifts affect treatment and control groups differently or when treatment effects vary across sites or years.

More broadly, this approach is not specific to the ITP. Interlaboratory testing is increasingly emphasized in aging research, as single-site, single-genotype findings have repeatedly failed to replicate across laboratories ^2^. Modeling between-cohort variance directly, with heteroscedasticity by sex or by group where warranted, could extend to other multi-site or multi-cohort preclinical aging studies that rely on a shared-control design, where a single unrecognized outlier cohort can otherwise propagate error across every treatment comparison made against it.

Limitations of the current framework are worth noting. With only three sites, the descriptive variation among fitted site means describes the ITP laboratories rather than laboratories in general. Kaplan–Meier estimation accounts for right censoring when constructing the cohort medians used in the historical reference analysis. The cohort-level model does not separately account for uncertainty in Kaplan–Meier median estimates, while the individual-level analyses use recorded ages without accounting for censoring. Censoring varied across sites, sexes, and years, particularly among UM males (Supplementary Figure 8), potentially influencing cohort means, variance estimates, and outlier classifications. Future survival-based analyses could help assess these effects. However, an exploratory analysis found no apparent positive relationship between censoring frequency and Kaplan–Meier median lifespan among UM male cohorts (data not shown), suggesting that their relatively high median lifespan is not simply explained by greater censoring. Although the prospective prediction intervals were constructed at a nominal 99% level, their empirical coverage was 95.0% across the 60 primary comparisons. This difference may reflect unusually extreme cohorts, imperfect calibration of the predictive model, or both, and warrants evaluation using additional independent cohort years. Finally, the framework identifies a cohort as statistically unusual, but cannot establish whether it reflects a procedural problem, environmental change, biology, or simply a rare realization of the normal process.

An unresolved question is how uncertainty in control-group variance should be incorporated into sample-size determination for lifespan studies. The historical heterogeneity observed here suggests that treating variance as known may understate uncertainty in power calculations. The implications of historical control heterogeneity for power analysis and sample-size determination warrant dedicated methodological investigation and are beyond the scope of the present study.

In the future, this framework should be applied prospectively to all newly completed control cohorts to identify those that are statistically unusual. Flagged cohorts should trigger prespecified sensitivity analyses of the corresponding treatment comparisons. Additionally, a complementary robustness check could be performed for treatment comparisons, especially when a control cohort is flagged as a potential outlier. Rather than comparing a candidate intervention only against its own parallel-year control cohort, the comparison could be iterated against a range of non-parallel-year historical control cohorts to test whether an apparent treatment effect depends on the particular control cohort used as reference or remains robust across the range of historical control variability characterized here. More formal approaches could also incorporate historical controls using power-prior or Bayesian borrowing methods^18;19^.

Control cohorts are the measuring instrument of an intervention study, and their variability sets the resolution of every comparison made against them. Eighteen years of parallel testing in the ITP show that control-cohort variability reflects substantial year-to-year variation, particularly in males, together with individual-level sampling variation and more modest systematic differences among the three testing sites. Applied prospectively, the framework provides a transparent basis for deciding when a control cohort warrants closer review before treatment effects are interpreted.

## Methods

### ITP control lifespan data

We analyzed control mice from the National Institute on Aging Interventions Testing Program at The Jackson Laboratory (TJL), the University of Michigan (UM), and the University of Texas Health Science Center at San Antonio (UT). The dataset comprised 10,757 control mice from 18 completed cohort years spanning C2004–C2022; no C2008 cohort was present. Sex, testing site, cohort year, recorded age, and survival status were retained for analysis. For Kaplan–Meier analyses, animals recorded as natural deaths were treated as events, whereas animals censored before natural death, including euthanized or terminated animals, were right-censored at their recorded age. Kaplan–Meier analyses incorporated survival status, whereas analyses of observed age used the recorded age directly.

### Descriptive analyses

Annual site-by-sex cohorts were summarized by mean observed age, standard deviation, and standard error for the descriptive table. Pooled survival curves and annual cohort-specific median survival were estimated by the Kaplan–Meier method. Figure 1A and Supplementary Figure 1 show annual cohort-specific Kaplan–Meier medians with 95% confidence intervals obtained by inversion of survival-curve confidence limits calculated using the log transformation. Temporal associations between cohort year and Kaplan–Meier median survival were examined using Pearson correlations separately for each site-by-sex combination, with *P* values adjusted across the six comparisons using the Benjamini–Hochberg method. The TJL-female analysis was additionally repeated after excluding C2022. Censoring frequency was examined descriptively by calculating the percentage of animals recorded as right-censored (Dead = 0) within each site *×* sex *×* year control cohort. Distributions of recorded age were also compared between censored animals and animals recorded as natural deaths (Dead = 1), stratified by site and sex. Density distributions were normalized separately within each censoring-status group.

### Variance partitioning

Individual-level observed age was modeled using linear mixed-effects models with sex, site, and their interaction as fixed effects. Site *×* year was modeled as a random effect with separate variance components for females and males using a diagonal covariance structure; testing site itself remained a fixed effect. A shared site *×* year variance and the sex-specific structure were compared using AIC and BIC because their fixed-effect structures were identical. Residual variance was allowed to differ by sex, and models with common versus sex-specific residual variance were compared by likelihood-ratio test. Sex *×* site marginal means and 99% confidence intervals were estimated using 5,000 wild-bootstrap resamples with Mammen auxiliary weights and HC2 leverage adjustment. Variability in an observed cohort mean was partitioned into systematic differences among fitted site means, sex-specific random site *×* year variance, and individual residual variance divided by the mean cohort size.

### Historical reference intervals

For the complete-record analysis, one Kaplan–Meier median lifespan was estimated for each site *×* sex *×* year cohort and modeled as a function of sex, with site as a random effect and sex-specific residual variance. The final reference model was fitted by restricted maximum likelihood. Sex-specific 99% prediction intervals for the estimated Kaplan–Meier median of a future comparable cohort were obtained from 100,000 simulations incorporating the fitted site and residual variation. Cohort-specific uncertainty in the Kaplan–Meier medians and uncertainty in the estimated model parameters were not modeled separately. To assess robustness, alternative models of Kaplan–Meier median lifespan included a linear year effect, site *×* sex interaction, site- and sex-specific linear temporal drift, or cohort size. Models containing fixed site effects were fitted using generalized least squares with sex-specific residual variances, generating prediction intervals conditional on testing site, whereas the remaining models retained a random site intercept. Sensitivity prediction intervals were calculated analytically from the fitted normal predictive distributions. Models were fitted using restricted maximum likelihood for prediction and maximum likelihood for model comparisons. Model fit was compared using AIC, with likelihood-ratio tests restricted to nested models with compatible covariance structures. Additional sensitivity analyses examined cohort mean observed age and raw median observed age rather than Kaplan–Meier median lifespan. The reference model was also separately refitted after excluding all cohorts from each of three candidate years (C2009, C2017, and C2022), evaluating each corresponding candidate against its independently estimated prediction interval.

### Prospective temporal validation

To approximate prospective use, the individual-level model was repeatedly refitted to cumulative historical windows, after which each fitted model was used to predict the mean lifespan of the subsequent site*×* sex cohort. For each window, 5,000 wild-bootstrap estimates of the fitted sex *×* site mean were generated. For each bootstrap estimate, a future sex-specific Site*×* Year effect was simulated from the fitted year-level variance and cohort-mean residual error was simulated using the fitted sex-specific residual variance divided by the observed size of the subsequent cohort. The 0.5th and 99.5th percentiles of the resulting simulated future cohort means formed the two-sided 99% prediction interval. Windows containing fewer than eight historical cohort years were retained for sensitivity analyses but excluded from the primary coverage summary; the eight-year minimum was retained as a pragmatic stability threshold because the average ratio of wild-bootstrap to model-based standard errors largely stabilized after approximately seven to eight historical cohort years. A sensitivity analysis additionally allowed a separate linear temporal slope for each site *×* sex combination by including the full interaction among sex, site, and centered cohort year as fixed effects. Cohort year was centered within each historical training window. The model retained the same sex-specific site *×* year random-effect structure and sex-specific residual variances as the base model. Prospective coverage and prediction-interval width were compared between the base and drift models over matching historical windows. Analyses were conducted in R version 4.5.1 (2025-06-13 ucrt).

### C2015 treatment sensitivity analysis

Because C2015 UM males were flagged under both prospective model specifications, we performed a secondary sensitivity analysis of the male interventions tested in C2015. Intervention assignments were identified from the original Rx variable. Each intervention was first compared with the parallel male controls using data from all three ITP sites and then reanalyzed after excluding all UM male animals, including both control and treatment-assigned mice. Survival differences were tested using log-rank tests stratified by testing site, consistent with the original C2015 analysis^16^. Kaplan–Meier median lifespan was calculated for each intervention and corresponding control group, and treatment effects were expressed as the percentage difference in median lifespan relative to control. Statistical significance was defined as P < 0.05.

## Conflict of interest statement

The authors declare no conflict of interest.

## Acknowledgments

We thank Richard A. Miller for promptly providing the most recent ITP lifespan data used in this analysis.

## Author contributions

LTM drafted the manuscript and performed the analyses. AL revised the initial manuscript and code, suggested additional analyses, and approved the final manuscript. JFN reviewed and approved the final manuscript. GAC conceptualized the study, revised the initial manuscript and code, and reviewed and approved the final manuscript.

## Notes

### Competing Interest Statement

The authors have declared no competing interest.

