## Supplementary Information for "A prediction-interval framework for identifying outlier control cohorts in lifespan studies"

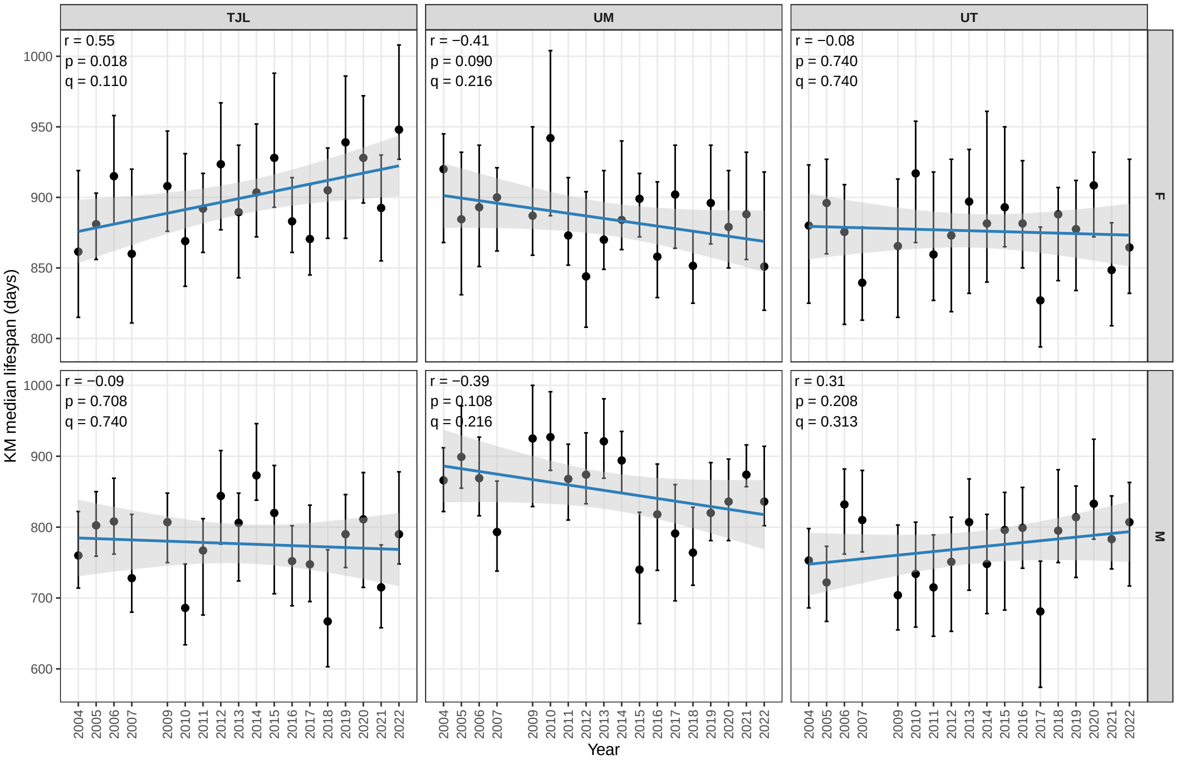


**Supplementary Figure 1. Temporal variation in control-cohort median lifespan by testing site and sex.** Annual Kaplan–Meier median lifespan is shown for female (top) and male (bottom) control cohorts at The Jackson Laboratory (TJL), University of Michigan (UM), and University of Texas Health Science Center at San Antonio (UT). Points represent the KM median lifespan within each site × sex × year cohort, and error bars show 95% confidence intervals obtained by inversion of the log-transformed Kaplan–Meier survival-curve confidence limits. Blue lines show linear fits with 95% confidence bands. Pearson correlation coefficients (r) and corresponding P values were calculated separately for each site × sex combination; q values were adjusted across the six comparisons using the Benjamini–Hochberg procedure.

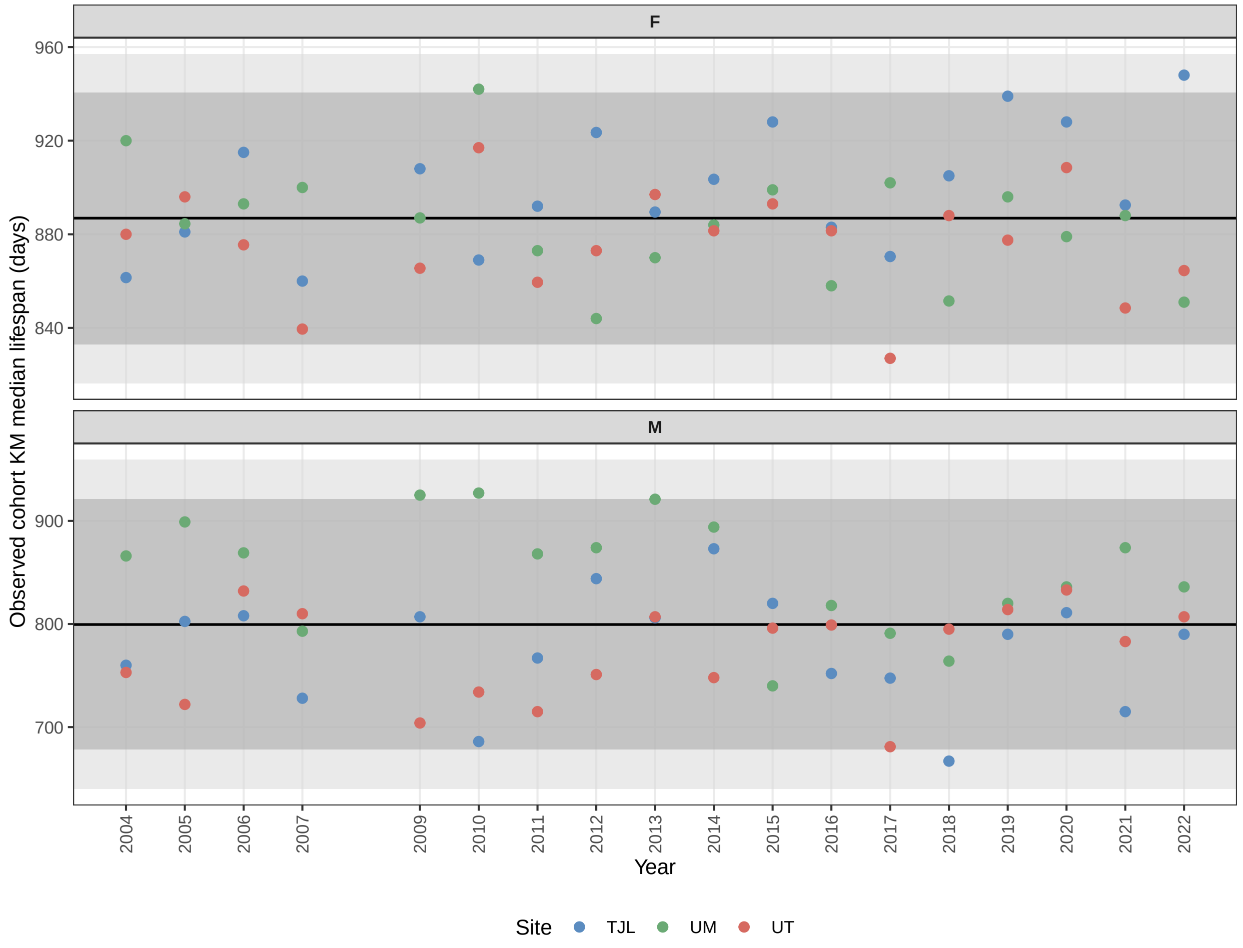


**Supplementary Figure 2. In-sample coverage of the complete-record cohort-median reference model.** Estimated Kaplan–Meier median lifespans for each historical site × sex × year control cohort are plotted against the sex-specific predictive ranges generated from the complete C2004–C2022 control record. Points are colored by testing site. The horizontal black line denotes the center of the predictive distribution, the darker gray region denotes the 95% prediction interval, and the lighter gray region denotes the 99% prediction interval. Panels are shown separately for females and males.

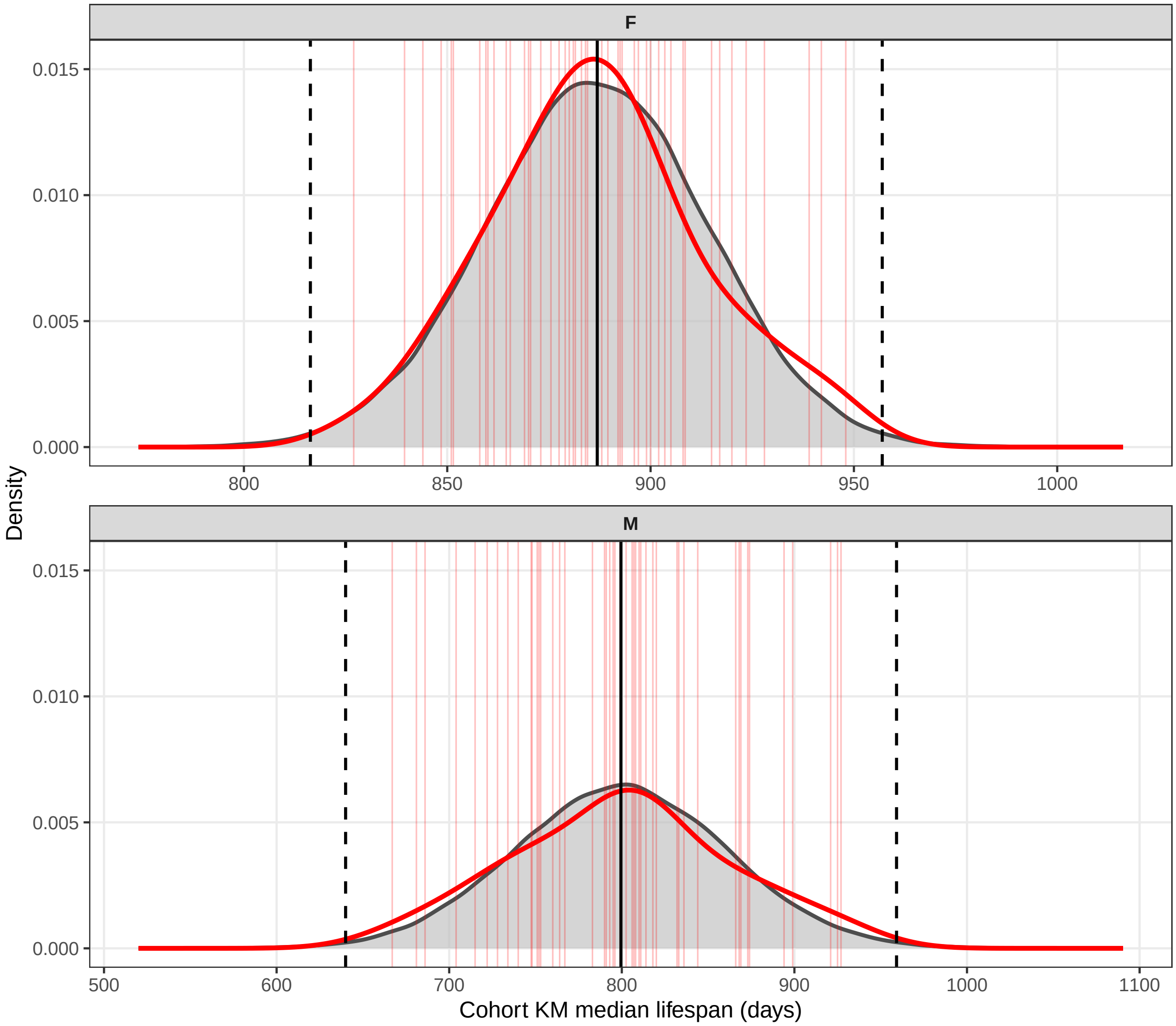


**Supplementary Figure 3. Agreement between observed and model-predicted distributions of control-cohort Kaplan–Meier median lifespan.** Gray distributions show simulated future cohort Kaplan–Meier medians from the complete-record reference model, whereas red curves show the empirical distributions of historical cohort Kaplan–Meier median estimates. Individual historical cohort Kaplan–Meier medians are indicated by faint red vertical lines. Solid black vertical lines indicate the model-predicted centers and dashed black lines indicate the sex-specific 99% prediction limits. Females and males are shown separately.

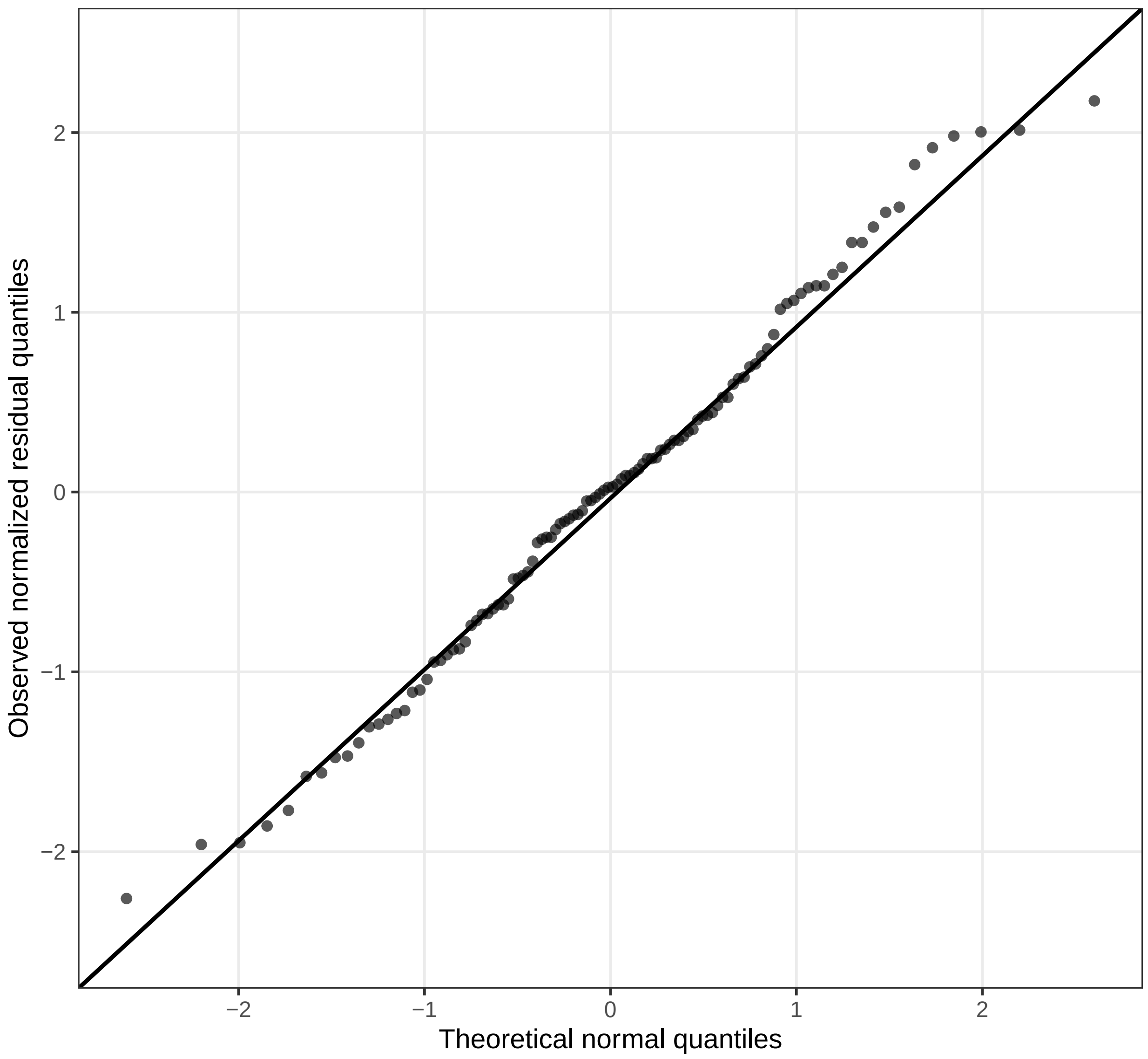


**Supplementary Figure 4. Residual diagnostic for the complete-record cohort-median reference model.** Quantile–quantile plot of normalized residuals from the linear mixed-effects model of cohort-specific Kaplan–Meier median lifespan. Points show observed residual quantiles plotted against theoretical normal quantiles; the reference line indicates the expected relationship under approximate normality.

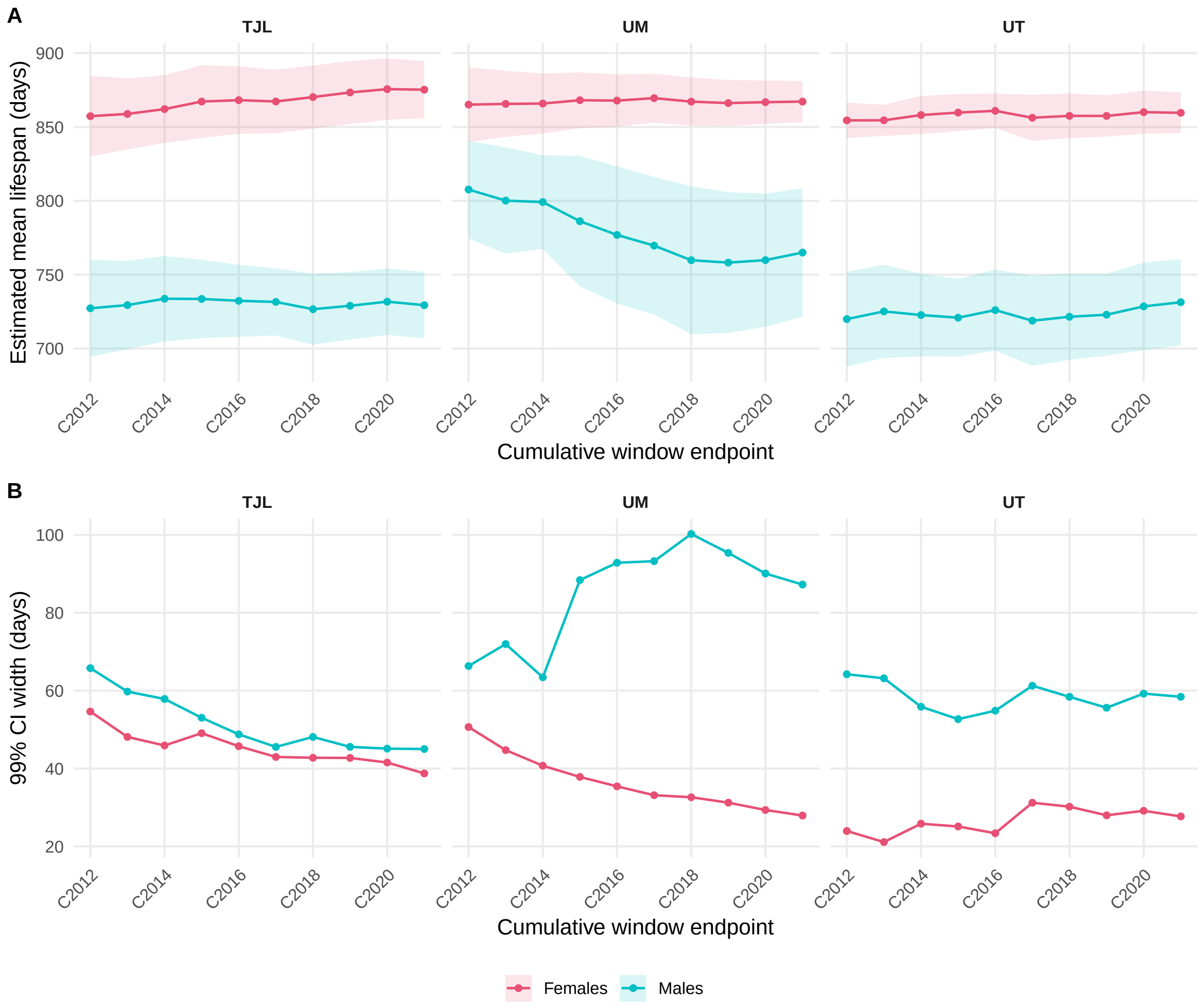


**Supplementary Figure 5. Stability of control-lifespan estimates as historical cohort data accumulate.** Models were refitted to cumulative historical windows ending at successive cohort years. A) Estimated mean lifespan for each site × sex combination; shaded regions show approximate 99% confidence intervals calculated using the standard errors from 5,000 wild-bootstrap resamples and a normal approximation. B) Width of the corresponding approximate 99% confidence intervals. Each point represents a model fitted using all historical cohorts available through the indicated endpoint. Only windows meeting the eight-year minimum used for the primary prospective analysis are shown.

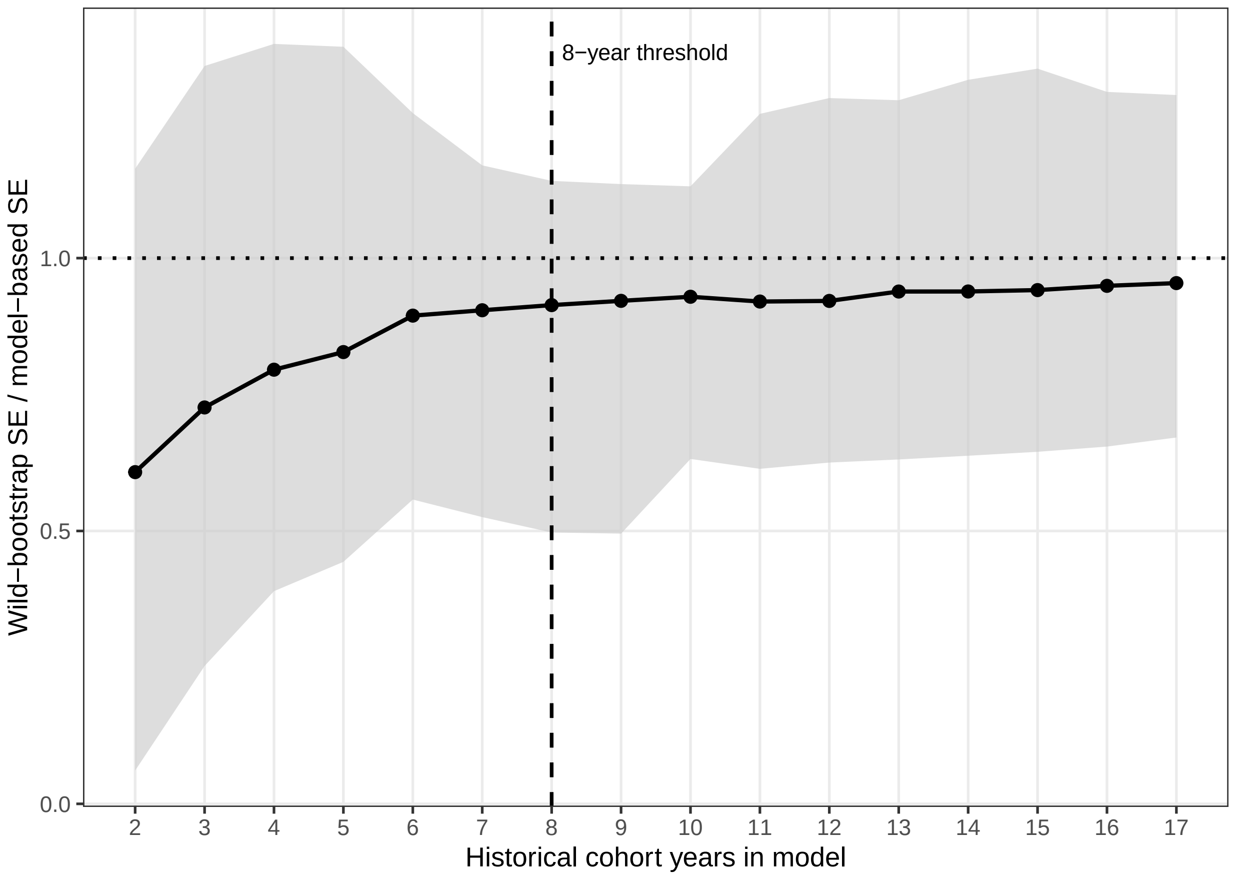


**Supplementary Figure 6. Descriptive diagnostic used to select the minimum historical-window length for prospective analysis.** Agreement between wild-bootstrap and model-based standard errors was evaluated as the number of historical cohort years increased. The y-axis shows the ratio of the wild-bootstrap standard error to the model-based standard error, with a value of 1 indicating exact agreement. The black line shows the mean ratio across site × sex estimates and the gray region shows the corresponding minimum-to-maximum range. The horizontal dotted line indicates a ratio of 1, and the vertical dashed line marks the eight-year reporting threshold used for the primary prospective analysis. The eight-year cutoff was retained as a pragmatic minimum because the average ratio had largely stabilized by approximately seven to eight historical cohort years and showed little systematic improvement in later windows.

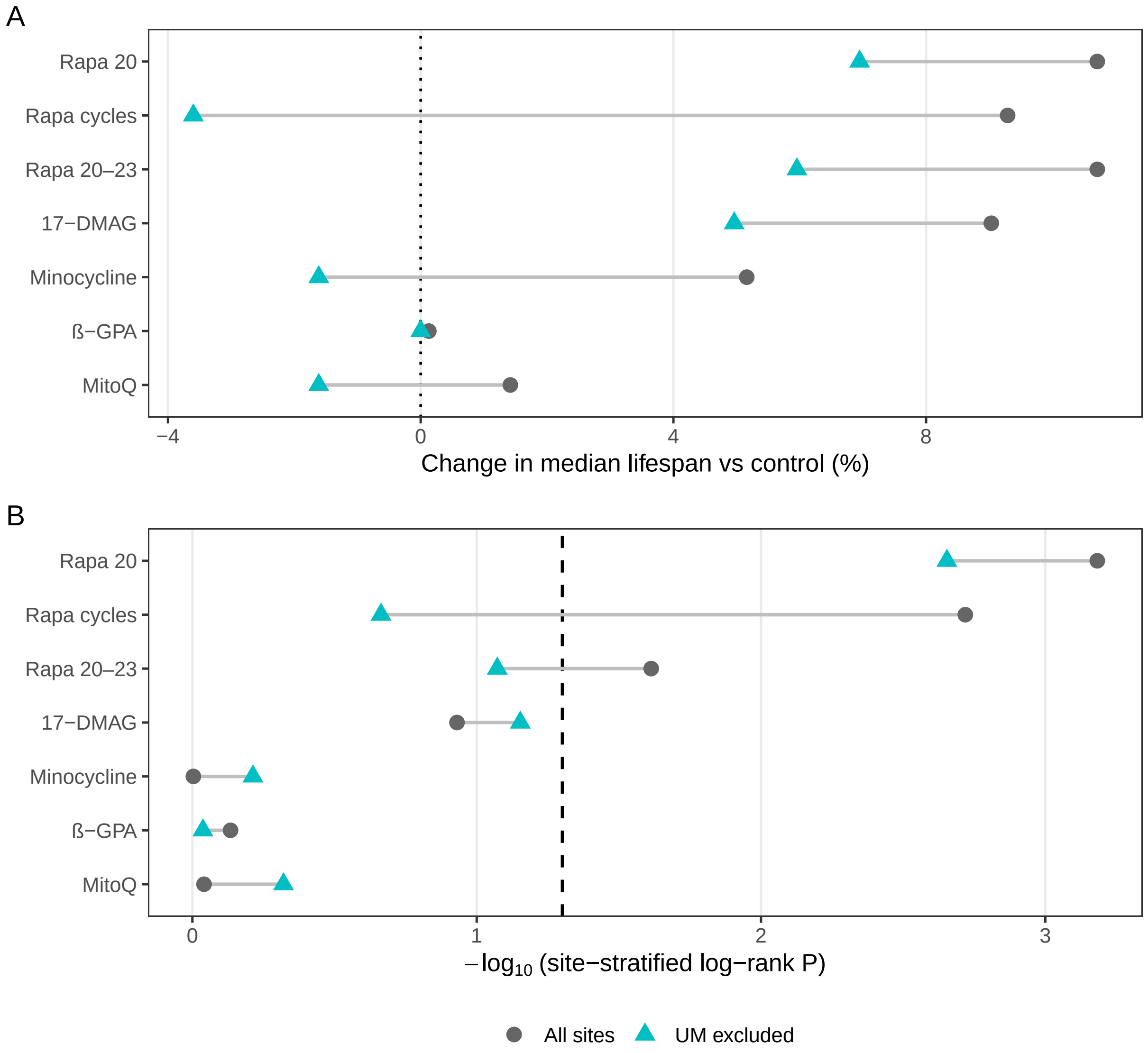


**Supplementary Figure 7. Sensitivity of C2015 male intervention effects to exclusion of the University of Michigan cohort.** C2015 male intervention effects were analyzed using all three ITP sites and after excluding all University of Michigan (UM) male animals, including both control and treatment-assigned mice. A) Percent change in Kaplan–Meier median lifespan relative to the corresponding control group. B) Site-stratified log-rank P values; the dashed line indicates $P = 0.05$. Exclusion of UM changed the nominal statistical significance of the Rapa cycles and Rapa 20–23 interventions, whereas Rapa 20 remained significant.

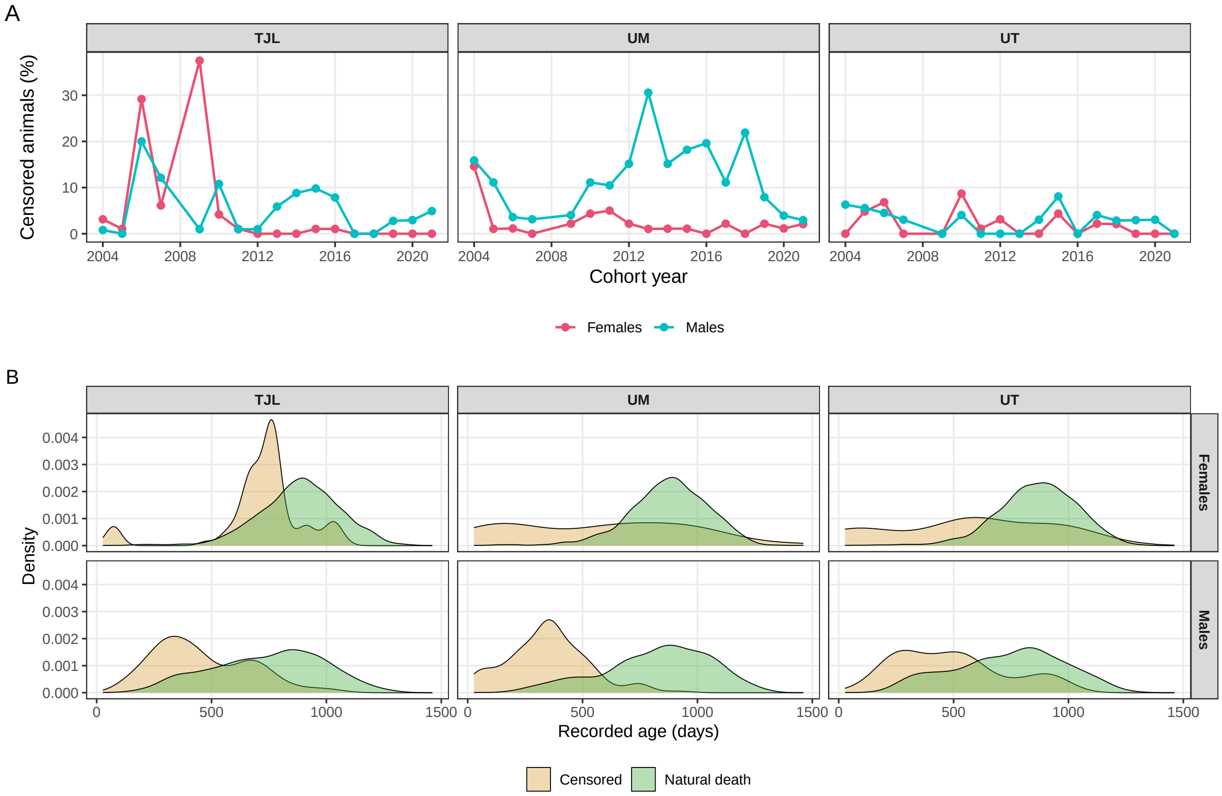


**Supplementary Figure 8. Frequency and timing of censoring among ITP control mice.** A) Percentage of control animals recorded as censored (Dead = 0) across cohort years, stratified by testing site and sex. B) Distributions of recorded ages among censored animals (Dead = 0) and animals recorded as natural deaths (Dead = 1), stratified by testing site and sex. Density distributions were normalized separately within each censoring-status group and therefore illustrate differences in the timing of recorded ages rather than the relative frequency of censoring.
